## supporting information for "ProTDyn: a foundation Protein language model for Thermodynamics and Dynamics generation"

### A Experiment details

#### A.1 Distributional similarity evaluation

**Radius of gyration.** The radius of gyration ($R_g$) is a fundamental descriptor of protein structure, quantifying the overall compactness of a molecule. It is defined as the root-mean-square distance of the constituent atoms from their common center of mass, thereby capturing how mass is distributed around the protein's centroid.

The radius of gyration is defined as

$$R_g = \sqrt{\frac{1}{N}\sum_{i=1}^{N} \|\mathbf{r}_i - \mathbf{r}_{\text{cm}}\|^2}, \tag{11}$$

where $N$ is the number of atoms, $\mathbf{r}_i$ is the position vector of the $i$-th atom, and $\mathbf{r}_{\text{cm}}$ is the position vector of the center of mass. This expression measures the spatial dispersion of the atomic positions relative to the center of mass. In this work, $R_g$ is computed using only the backbone atoms.

**Root Mean Square Distance (RMSD).** The root mean square distance (RMSD) is a standard metric in structural biology for quantifying the similarity between two protein conformations. RMSD is computed by superimposing the two structures and calculating the square root of the average squared distance between corresponding atoms:

$$\text{RMSD} = \min_{T_g \in \text{SE}(3)} \sqrt{\frac{1}{N}\sum_{i=1}^{N} \left\|T_g(\mathbf{r}_i) - \mathbf{r}_i^{\text{ref}}\right\|^2}, \tag{12}$$

where $N$ is the number of atoms, $\mathbf{r}_i$ are the atomic coordinates of the structure under comparison, $\mathbf{r}_i^{\text{ref}}$ are the coordinates of the reference structure, and $T_g \in \text{SE}(3)$ denotes the optimal rigid-body transformation (rotation and translation) aligning the two structures. Lower RMSD values indicate greater structural similarity.

**Time-lagged Independent Component Analysis (TICA).** TICA is a linear method for extracting the slowest dynamical modes from time-series data, widely used in molecular dynamics. Unlike PCA, which finds directions of largest variance, TICA identifies directions with maximal autocorrelation at lag time $\tau$. It solves the generalized eigenvalue problem

$$C_\tau r_i = C_0 \lambda_i r_i, \tag{13}$$

where $C_0$ is the covariance matrix, $C_\tau$ is the time-lagged covariance, and $\lambda_i$ are the time-autocorrelations. The resulting components capture the slow collective motions that dominate long-timescale protein dynamics. In this project, we choose the feature as protein backbone torsion angles, and extract the top 2 TIC components.

#### A.2 Dynamical contents evaluation

**Autocorrelation** We define the autocorrelation of each TICA component as

$$\mathbb{E}\big[(y_t - \mu)(y_{t+\delta t} - \mu)\big]/\sigma^2 \tag{14}$$

where $\mu, \sigma$ are computed from the reference MD simulation trajectory. We conduct TICA on lag times $\delta t = [10, 20, \ldots, 800]$ ns.

**Markov state models** Markov state models (MSMs) provide a statistical framework for describing the long-timescale dynamics of biomolecules by coarse-graining the continuous conformational space into a finite set of metastable states. The dynamics are modeled as a discrete-time Markov chain, where the probability of transitioning between states depends only on the current state and a chosen lag time $\tau$. Formally, the state-to-state transition probabilities are encoded in a transition matrix $T(\tau)$:

$$p_j(t+\tau) = \sum_i p_i(t)\, T_{ij}(\tau), \tag{15}$$

where $p_i(t)$ is the probability of being in state $i$ at time $t$, and $T_{ij}(\tau)$ is the probability of transitioning from state $i$ to $j$ over lag time $\tau$. At equilibrium, MSMs satisfy the stationary distribution condition

$$\boldsymbol{\pi} = \boldsymbol{\pi} T(\tau), \quad \sum_i \pi_i = 1, \tag{16}$$

where $\pi_i$ denotes the equilibrium probability of state $i$.

We follow previous works Jing et al. (2024b); Raja et al. (2025) and build MSM using Deeptime Hoffmann et al. (2021). We first represent protein systems with backbone torsion angles and run TICA to obtain the top 2 Time Independent Component dimensions. We then perform k-means clustering of the reference MD simulations into 10 clusters. We then fit a MSM with a lag time of 10 ns. This gives us the transition probability matrix $T$. The stationary distribution $\pi$ can be easily obtained as the left eigenvector of the transition matrix with eigenvalue 1. We use the same settings to construct MSM for reference MD simulations, dynamic trajectories generated by ProTDyn, and different portions of reference MD simulations.

We evaluate the following two metrics:

- **Stationary distribution Jenson-Shannon divergence** We evaluate the stationary distribution of each MSM state for reference and comparison trajectories, and compute the JSD between the categorical distributions.
- **Transition probability Jensen–Shannon divergence** We evaluate the transition probability matrix between MSM states for both reference and comparison trajectories, and compute the JSD between the corresponding transition probability distributions.
- **Trajectory negative log likelihood** We use the stationary distribution of reference MSM to generate 1000 initial states: $s_0 \sim \pi_{\text{ref}}$, and then propagate a dynamic path of length $L = 20$ from each iniitial state for each comparison MSM using its transition proability matrxi: $s_{t+1} \sim T_{s_t}$. The probability of a path can thus be written as $P(s_1, ..., s_L) = -\frac{1}{L}\sum_{t=1}^{L} \log T_{\text{ref}_{s_t, s_{t+1}}}$. We report the mean negative log likelihood over all trajectories.

### B Additional experiment results

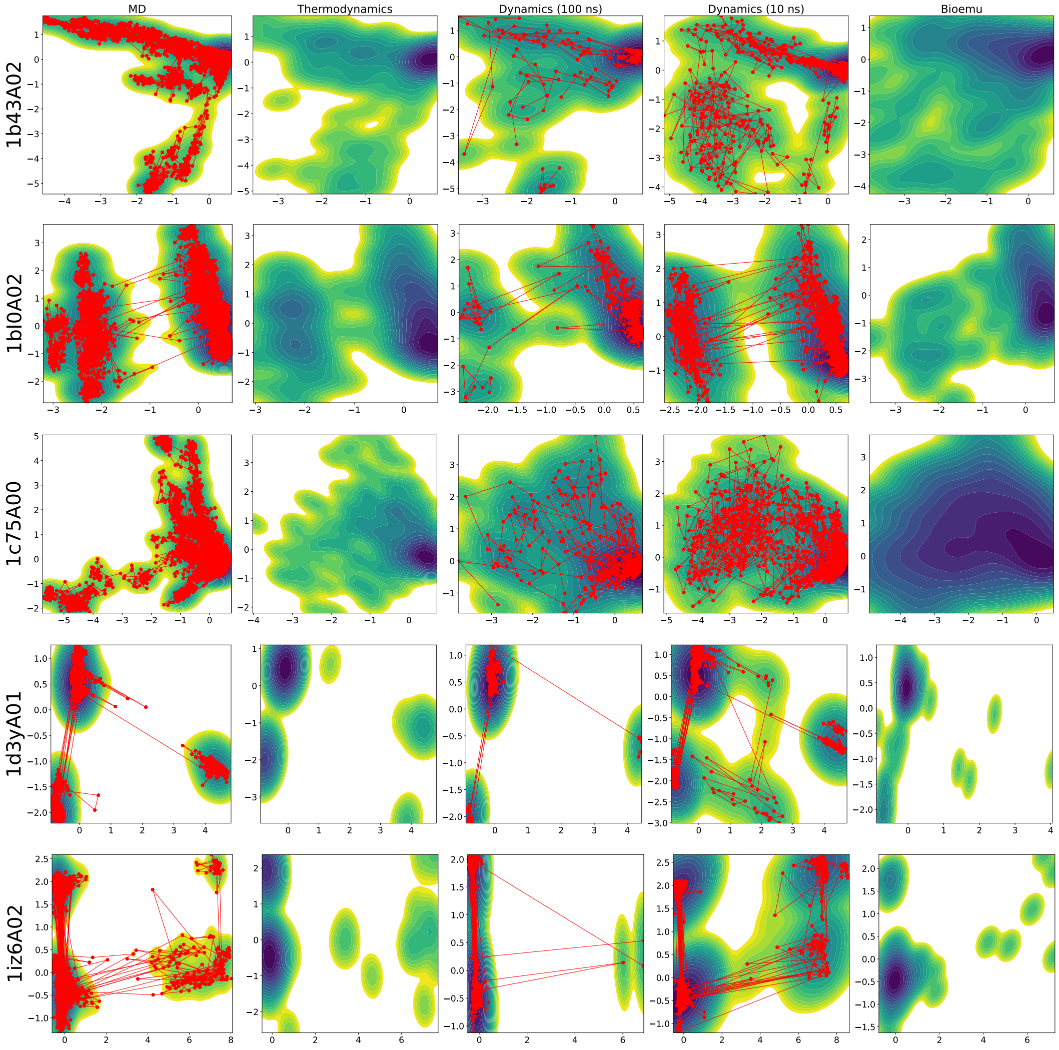


Figure 5: Additional results on 5 test CATH1 proteins: free energy surface along the top two TICA components and dynamic transition pathways. Note that the thermodynamics module of ProTDyn and baseline model Bioemu are both i.i.d sampler and thus does not have a dynamic trajectory.

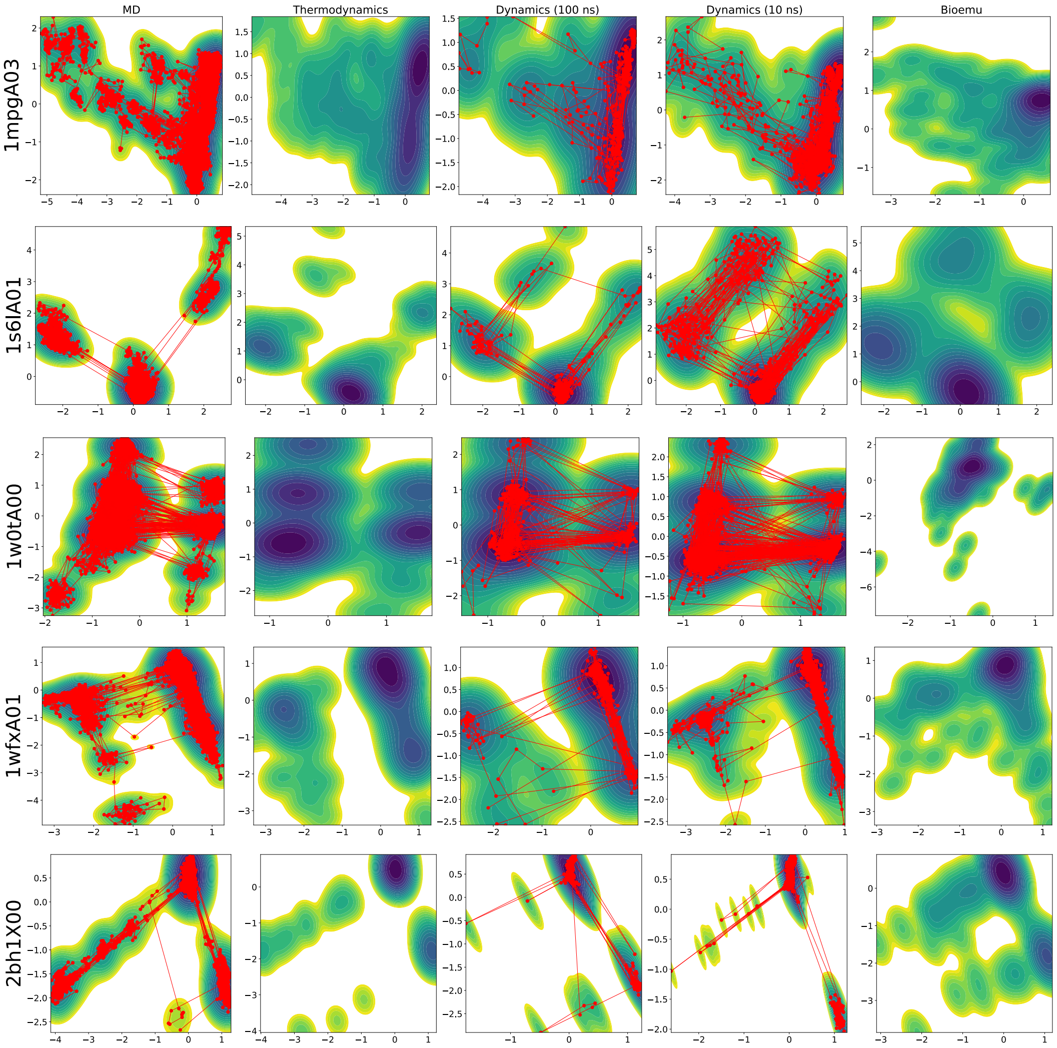


Figure 6: Additional results on another 5 test CATH1 proteins.

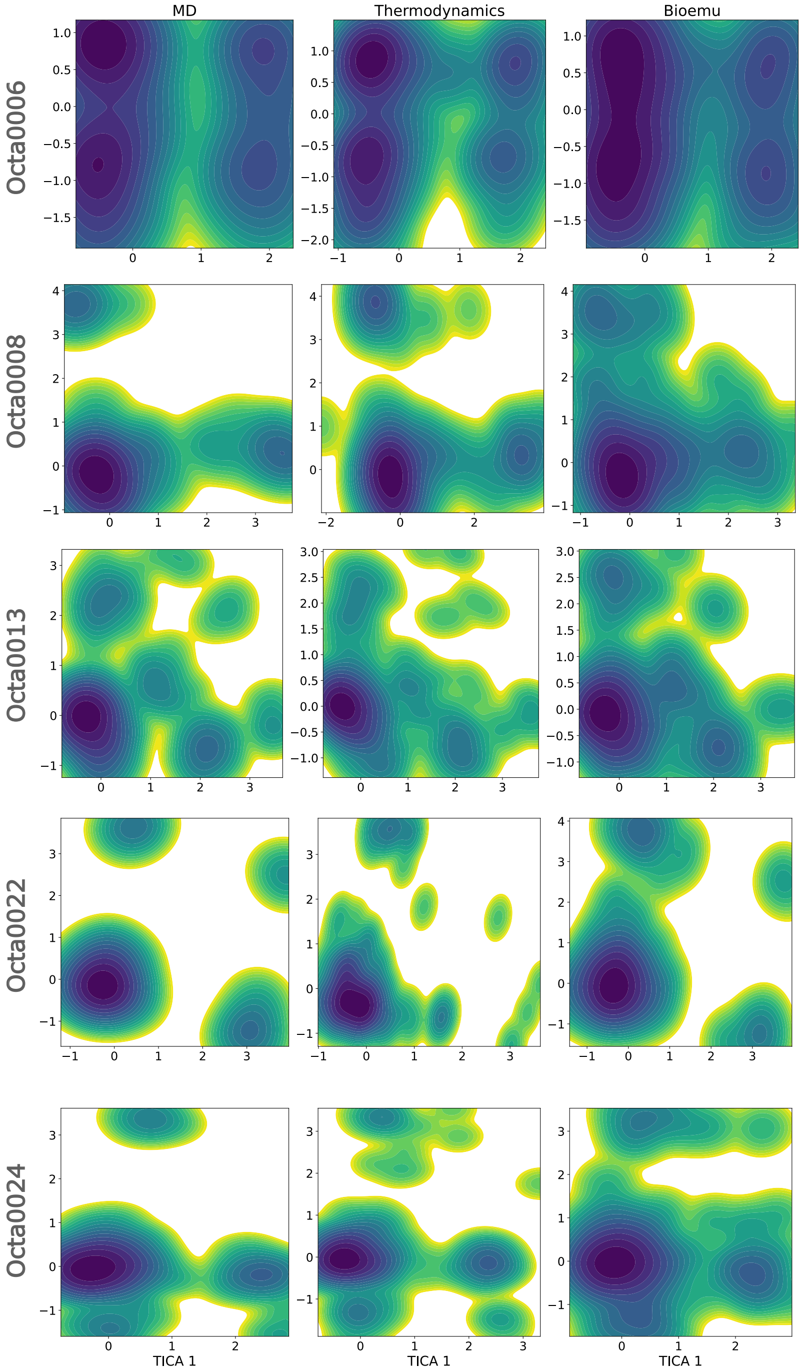


Figure 7: Additional results on 5 test Octapeptides: free energy surface along the top two TICA components.

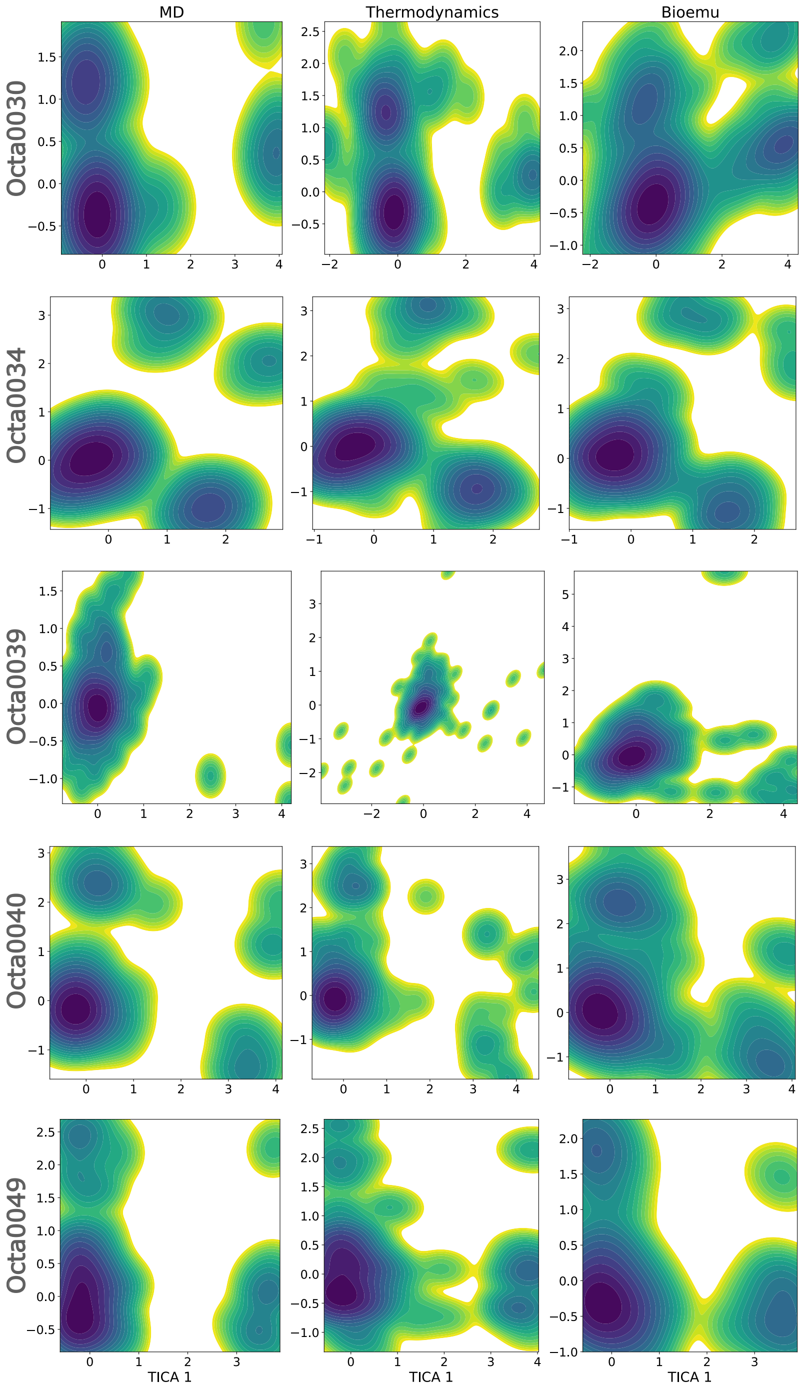


Figure 8: Additional results on another 5 test Octapeptides: free energy surface along the top two TICA components.
